# Reproductive longevity predicts mutation rates in primates

**DOI:** 10.1101/327627

**Authors:** Gregg W.C. Thomas, Richard J. Wang, Arthi Puri, R. Alan Harris, Muthuswamy Raveendran, Daniel Hughes, Shwetha Murali, Lawrence Williams, Harsha Doddapaneni, Donna Muzny, Richard Gibbs, Christian Abee, Mary R. Galinski, Kim C. Worley, Jeffrey Rogers, Predrag Radivojac, Matthew W. Hahn

## Abstract

Mutation rates vary between species across several orders of magnitude, with larger organisms having the highest per-generation mutation rates. Hypotheses for this pattern typically invoke physiological or population-genetic constraints imposed on the molecular machinery preventing mutations^1^. However, continuing germline cell division in multicellular eukaryotes means that organisms with longer generation times and of larger size will leave more mutations to their offspring simply as a by-product of their increased lifespan^2,3^. Here, we deeply sequence the genomes of 30 owl monkeys (*Aotus nancymaae*) from 6 multi-generation pedigrees to demonstrate that paternal age is the major factor determining the number of *de novo* mutations in this species. We find that owl monkeys have an average mutation rate of 0.81 × 10^−8^ per site per generation, roughly 32% lower than the estimate in humans. Based on a simple model of reproductive longevity that does not require any changes to the mutational machinery, we show that this is the expected mutation rate in owl monkeys. We further demonstrate that our model predicts species-specific mutation rates in other primates, including study-specific mutation rates in humans based on the average paternal age. Our results suggest that variation in life history traits alone can explain variation in the per-generation mutation rate among primates, and perhaps among a wide range of multicellular organisms.

---

The rate at which new mutations arise is a key parameter of life on Earth, contributing to both individual disease risk and the evolution of novel traits. The mutation rate per generation varies among taxa, from as low as 1×10^−10^ per base in Archaea to more than 1×10^−8^ in mammals^1^. Two classes of models have been proposed to explain this variation. In one, the physiological and biochemical costs of increased fidelity during DNA replication limit the minimum mutation rate achievable^4,5^. Selection for faster replication in smaller organisms constrains the accuracy with which the cellular machinery can copy DNA, resulting in an inverse relationship between body size and mutation rate. Alternatively, a population-genetic model invokes the limits to natural selection in organisms with smaller population sizes^6–8^. This model posits a higher rate of mutation in larger organisms because of their generally smaller population size^9^.

One difficulty in teasing apart the forces driving the evolution of the mutation rate among multicellular organisms is the fact that lifespan varies as much as the per-generation mutation rate. In multicellular organisms, the number of mutations passed on to offspring in a single generation is a combination of the errors made in each round of germline replication and the accumulation of unrepaired DNA damage. One hundred years after the first observation of increased disease incidence in the children of older parents^2,10^, whole-genome sequencing in humans revealed the precise contribution of parental age to the number of *de novo* mutations in their offspring^3,11–17^. In particular, the number of mutations passed on to the next generation is largely dependent on the age of the father^3^, though there is a non-negligible contribution from the age of the mother^12–15,17^. This is a consequence of the fact that after a set number of germline mitoses during development in both males and females, the male germline resumes cell division at puberty^18,19^. A similar effect of paternal age has been found in chimpanzees^20^, suggesting that the age of reproduction may generally be an important determinant of the per-generation mutation rate.

Studying closely related primates offers a unique opportunity to examine the role that life history traits—such as age of puberty and average generation time—may play in determining mutation rates. We sequenced the genomes of 30 owl monkeys (*Aotus nancymaae*) within 6 multi-generation pedigrees (Fig 1a; Table S1) in order to estimate the effect of parental age on the mutation rate. Owl monkeys reach sexual maturity at ~1 year of age^21^ and can live up to 20 years in captivity^22^. Our sample includes individuals conceived by sires ranging from 3-13 years old and dams ranging from 3-12 years old, with an average age of 6.64 and 6.53 for sires and dams, respectively (Table S1). These ages are comparable to those observed in the wild, as owl monkeys are solitary for some time before joining a mating group at around age four^23^. The genomes of all parents and offspring were sequenced to an average of 37X coverage (range: 35X-38X) using paired-end Illumina reads. Sequencing multi-generation pedigrees allows us to determine whether *de novo* mutations arose in either sires or dams, as well as to validate mutations transmitted to the next generation.

**Figure 1:**
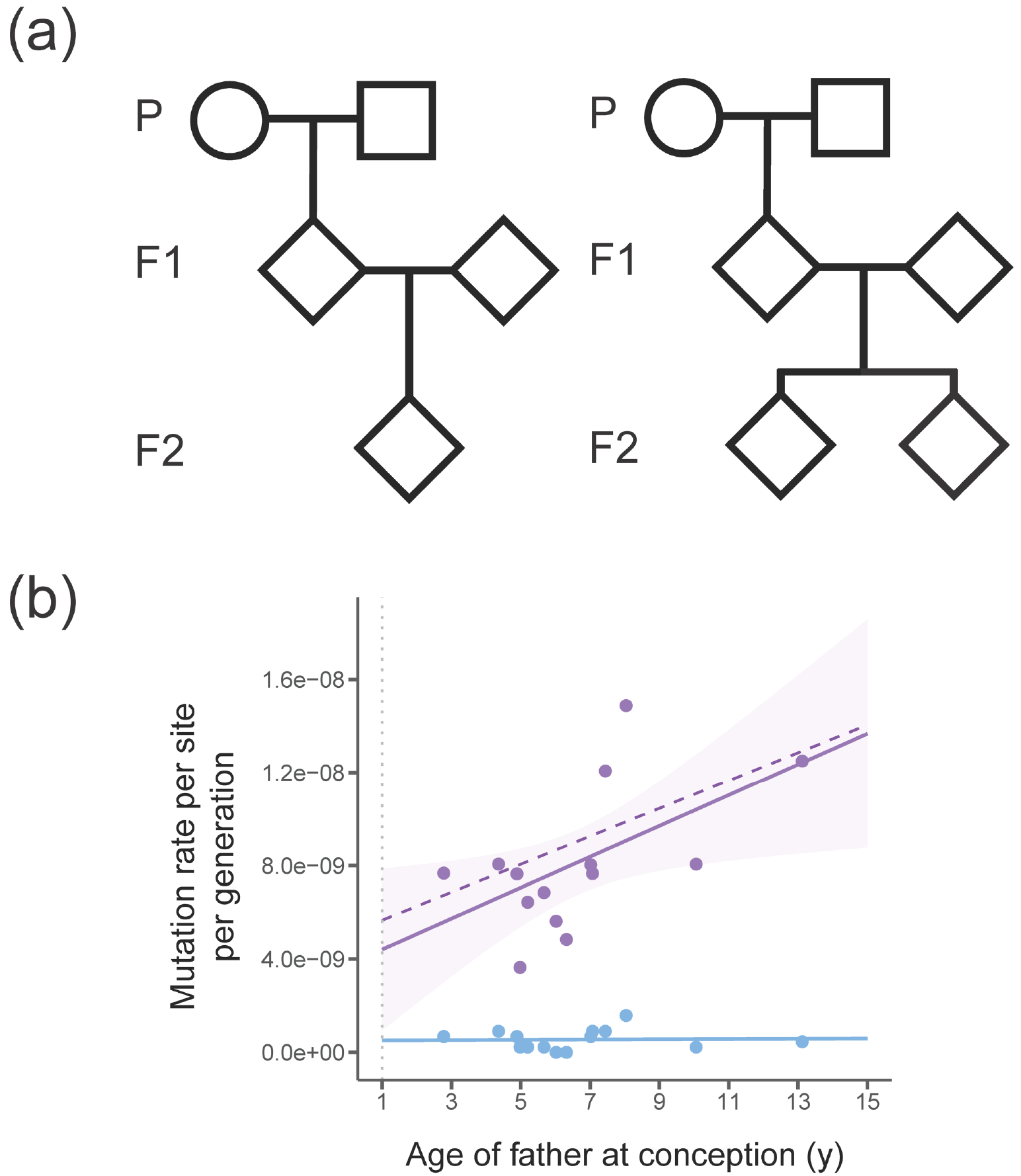
Pedigree structures and mutation rates in owl monkeys. **a,** We used six multi-generation pedigrees in these two formats. Four families have a single F2 offspring (left) while two families have two F2 offspring (right). In total, 14 independent trios can be constructed from these pedigrees. **b,** Mutation rate estimates from the 14 owl monkey trios (purple points). A simple linear regression has been fit to these points (solid purple line) to show that the number of mutations increases with the father’s age. Our model of reproductive longevity (dashed purple line) is not significantly different from the fit of the linear regression. The rate of non-replicative mutations, such as those that occur at CpG sites (blue dots), are not correlated with reproductive longevity (blue line). The dotted vertical grey line indicates expected age of puberty.

We observe 283 *de* novo mutations across 14 trios (Table S2) and estimate an average mutation rate for owl monkeys of 0.81 × 10^−8^ per site per generation (Table S3). In addition to stringent quality filters (see Methods), the average transmission frequency of *de novo* mutations passed from F1 individuals to F2 individuals across families was 0.502, giving us high confidence in our final set of mutations. As in humans, we find a strong association between paternal age and the number of *de novo* mutations (Fig 1b), with 2.92 additional mutations accumulating per year (*R^2^*=0.25, d.f.=12, *P*=0.040). Also as expected, we find no effect of age on CpG mutations (Fig 1b, blue points and line), as these are not associated with replication errors. We were able to assign phase to 105 of the 283 *de novo* mutations via transmission to the third generation in our pedigrees (Table S2). We find that 71 of these 105 phased mutations are paternal, with the number of mutations passed on increasing with the age of the father (*R^2^*=0.58, d.f.=4, *P*=0.048). We did not find an increasing number of mutations with maternal age (*R^2^*=0.07, d.f.=4, *P*=0.307). This is the first direct observation of the paternal age effect outside of apes.

Inspection of the types of mutations found in the genomes of owl monkeys shows a transition:transversion (Ts:Tv) ratio of 1.97. This is in close agreement with the observed human Ts:Tv ratio of 2.10 (ref. 3). In fact, the overall mutational spectrum between humans, chimpanzees, and owl monkeys appears almost identical, with the only difference being a slightly higher proportion of A→T mutations in owl monkeys (Fig 2). We also observe that 26.8% of mutations in owl monkeys occur at CpG sites, with CpG sites having a much higher Ts:Tv ratio (4.67), similar to observations in humans^3,14^. Multinucleotide mutations (MNMs) are mutations that occur in close proximity to one another (<20 bp apart), likely caused by a single mutational event^24^. Here, we find 6 MNMs consisting of two mutations each, indicating that 2.1% of *de novo* mutations in owl monkeys are the result of MNMs (Table S2). This fraction is also in agreement with that observed within humans^14,24^.

**Figure 2:**
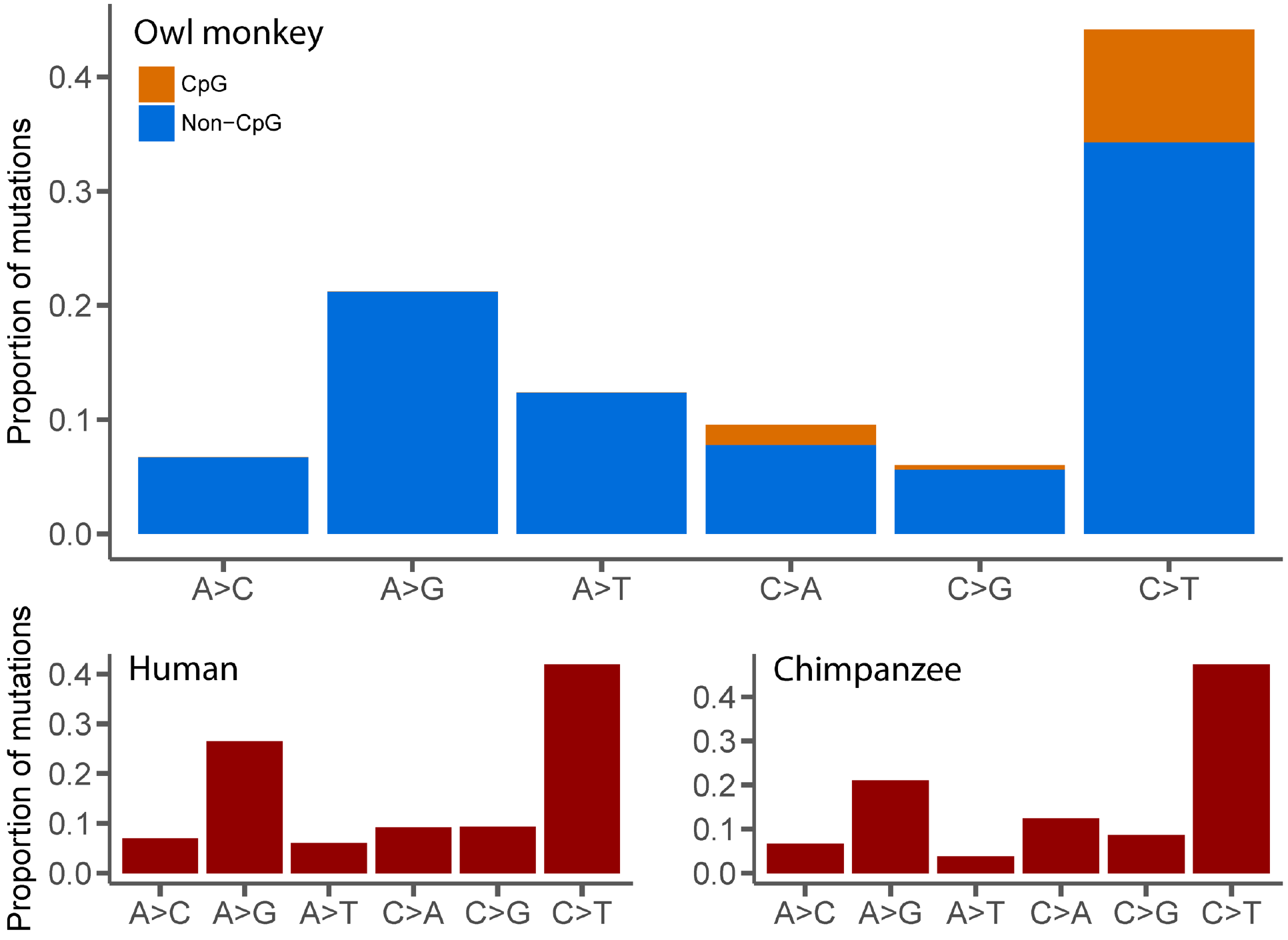
Comparison of mutational spectra from owl monkeys, humans, and chimpanzees. There is a slight but significant difference in the frequency of A→T mutations between owl monkeys and humans (*χ*^2^= 25.7, d.f. = 4, *P*<0.05), but otherwise no difference between mutational spectra for these three species. Human data were averaged across four studies (see Table S4 for references) and chimpanzee data was extrapolated from Figure 3A in Venn et al.^20^. Mutation categories include their reverse complement.

The mutation rate we observe in owl monkeys is 32.5% lower than the average human estimate of 1.2 × 10^−8^ mutations per site per generation^3,25^. While traditional models of mutation rate evolution invoke changes to the underlying replication machinery as the main cause of such differences, we asked whether a shift in reproductive timing could explain the lower rate in owl monkeys. The effects of paternal age on per-generation mutation rates have previously been modeled by combining estimates of the rate of mutation from different life stages^19,25,26^. The germline in males and females undergo a fixed number of divisions before birth, but the male germline continues dividing upon reaching sexual maturity. This phenomenon suggests that the length of time between puberty and the conception of offspring in an individual—which we define here as the *reproductive longevity* of males—plays a key role in determining the number of mutations passed on to the next generation. While paternal age is sufficient for predicting mutation rates within a species^3,11–17^, the concept of reproductive longevity makes it possible to predict mutation rates between species with varying ages of puberty. We modeled the owl monkey mutation rate as a linear combination of the mutations accumulated as a result of a constant number of germline divisions *in utero* and those accumulated during continued germline divisions post-puberty. The rate of mutation in these two stages were estimated from human studies, while sexual maturity was set at 1 year of age (Methods).

Our minimal model provides an excellent fit to the observed owl monkey data (Fig 1b, dashed line). In fact, a linear regression of the observed number of mutations with paternal age at conception is not significantly better than the fit provided by our model (*F*=0.996, d.f.=13, *P*=0.994). The main determinant of the mutation rate is reproductive longevity in sires, which determines the number of mitotic germline divisions before spermatogenesis. For instance, a 13-year-old owl monkey male (who reached sexual maturity at 1) will have the same reproductive longevity as a 25-year-old human male (who reached sexual maturity at 13). Our model therefore predicts the same estimated mutation rate if *de novo* mutations are sampled from offspring of these individuals, and this is what is observed (Fig 1b). Because reproductive longevity reflects replicative mutations, we observe no effect of father’s age on non-replicative mutations, such as those found at CpG sites (Fig 1b, blue).

Given the fit of our model to owl monkey data, we calculated the expected mutation rates as a function of age for other primates, accounting for changes in the time to sexual maturity in each species. A model of reproductive longevity provides a good fit to the data from primate species for which direct mutation rate estimates are available (Fig 3; Table S4). Our model explains why chimpanzees and humans have very similar per-generation mutation rates despite differences in average generation time: the earlier time to sexual maturity in chimpanzees causes reproductive longevity to be the same in both species. The model also accurately predicts estimated mutation rates reported from various studies in humans where sampled parents were of different average age (Fig 3). Much of the variation in reported mutation rates in human studies is due to differences in the average reproductive longevity of sampled individuals (*R^2^*=0.54, d.f.=7, *P*=0.01). Variation in the age of reproduction across pedigrees will affect inferences regarding genetic variation in the mutation rate, as consistent differences in these ages may incorrectly be interpreted as heritable differences in this trait.

**Figure 3:**
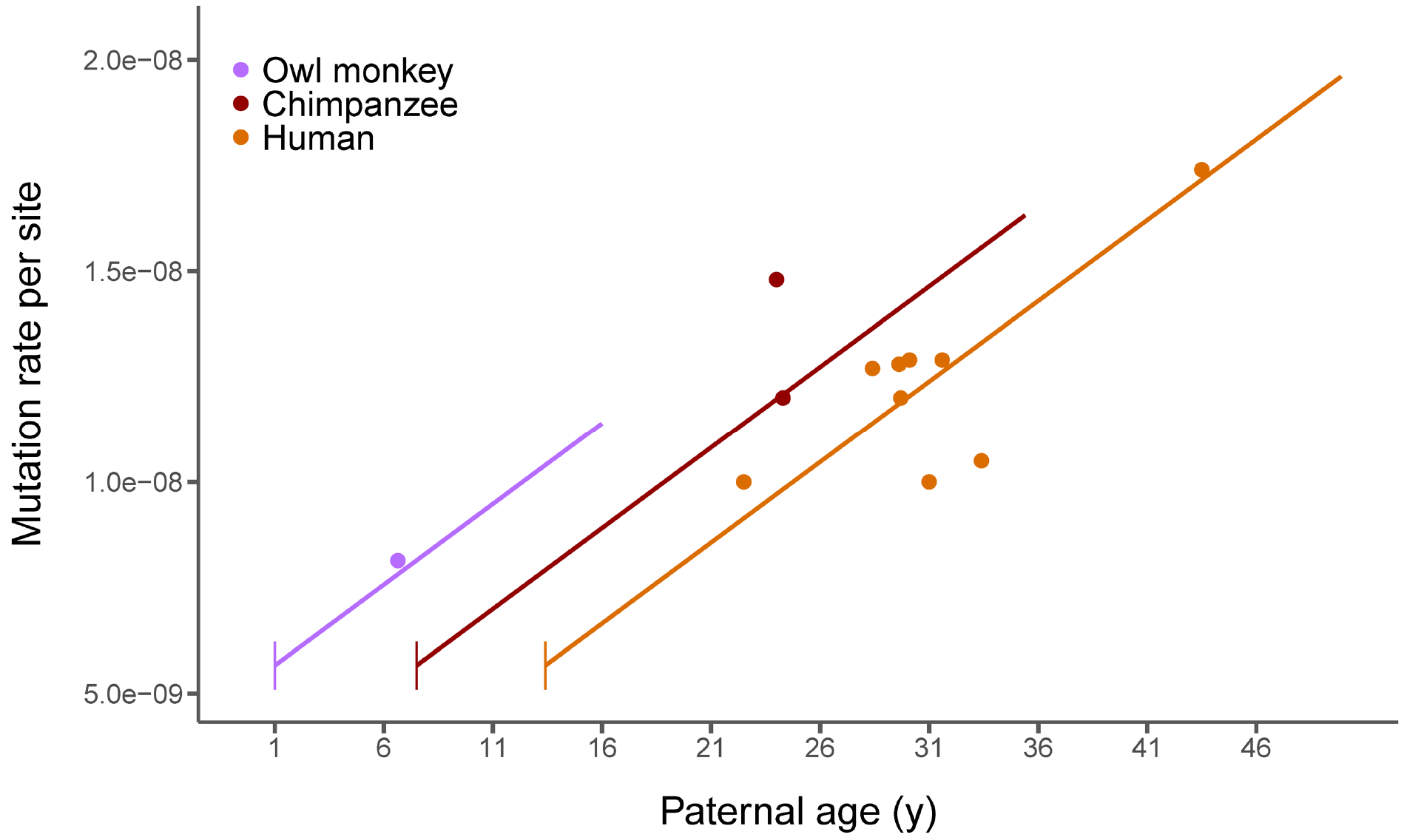
A model of reproductive longevity fits estimated primate mutation rates. Humans, chimpanzees, and owl monkeys are the only primates that currently have high-quality estimates of mutation rates via pedigree sequencing. Here we plot the average rate from each published study (points; see Table S4 for references). Predictions from our model of reproductive longevity (equations 3-8 in Methods) using human mutational parameters—varying only life history traits—are also shown (lines). Vertical line segments represent the age of puberty for each species.

The association between mutation rates and reproductive longevity implies that changes in life history traits rather than changes to the mutational machinery are responsible for the evolution of these rates. Species that have evolved greater reproductive longevity will have a higher mutation rate per generation without any underlying change to the replication, repair, or proofreading proteins. The similarities between the mutational spectra of humans, chimpanzees, and owl monkeys (Fig 2) are further evidence that the molecular mechanisms responsible for mutation have not changed between these species. Many differences in the details of germline cell division may exist between these primates, but these differences do not appear to affect either pre-birth or post-puberty mutation accumulation. The underlying consistency of mutation rates must also be reconciled with variation in the long-term substitution rate among primates^25–28^, as mutation rates are mechanistically tied to substitution rates. Nevertheless, the close fit between the observed and expected mutation rates suggests that reproductive longevity is the major determinant of variation in mutation rates.

Studies of mutation rate evolution will continue to accumulate across the tree of life as sequencing costs continue to plummet. In order to understand the forces affecting this important evolutionary parameter, future studies must recognize that the mutation rate is a function-valued trait: it is a function of reproductive longevity and other life history traits. Evidence from other species—for instance, arthropods^29^ and long-lived plants^30^—suggests that reproductive longevity affects the mutation rate in many taxa, though the details of germline cell division will differ among lineages. If such a pattern holds widely in multicellular organisms, the effect of variation in life history traits should provoke a reexamination of the causes underlying the correlation between body size and the per-generation mutation rate. At the very least, the null model for changes in the per-generation mutation rate must include reproductive longevity.

## Acknowledgements

This research was supported by the Precision Health Initiative at Indiana University. This research was also supported in part by Federal funds from the U.S. National Institute of Allergy and Infectious Diseases, National Institutes of Health, Department of Health and Human Services under contract # HHSN272201200031C, which supported the Malaria Host-Pathogen Interaction Center (MaHPIC).

## Author contributions

G.W.C.T., M.W.H., and P.R. conceived analyses; G.W.C.T., R.J.W., and A.P. performed analyses; M.W.H. supervised analyses; C.A. directs and supervises owl monkey colony; L.W. manages owl monkey colony and assisted with sample acquisition and meta-data; M.R. managed the sequencing process; M.R. and R.A.H called SNPs; R.A.H., D.H., and S.M., assembled and analyzed the reference genome; H.D. supervised library production and tech development; D.M. directs operations of sequencing and developed pipeline operations; R.G. directs the sequencing center and supervises sequencing operations; M.R.G. provided project rationale; K.C.W. and J.R. planned and supervised the genome assembly; M.W.H., P.R., and J.R. designed the project.

## Methods

### Sampling and sequencing

Thirty owl monkeys (*Aotus nancymaae*) were selected for genome sequencing from the Owl Monkey Breeding and Research Resource at the Keeling Center based on available pedigrees, aiming for a spread of parental ages (Table S1). Blood samples were taken from the femoral vein of unanesthetized animals. The animals were manually restrained in a supine position with one care staff holding the animal while another takes the sample, under approved IACUC protocols.

Genomic DNA isolated from the blood samples was used to perform whole genome sequencing. We generated standard PCR-free Illumina paired-end sequencing libraries. Libraries were prepared using KAPA Hyper PCR-free library reagents (KK8505, KAPA Biosystems Inc.) in Beckman robotic workstations (Biomek FX and FXp models). We sheared total genomic DNA (500 ng) into fragments of approximately 200-600 bp in a Covaris E220 system (96 well format) followed by purification of the fragmented DNA using AMPure XP beads. A double size selection step was employed, with different ratios of AMPure XP beads, to select a narrow size band of sheared DNA molecules for library preparation. DNA end-repair and 3’-adenylation were then performed in the same reaction followed by ligation of the barcoded adaptors to create PCR-Free libraries, and the library run on the Fragment Analyzer (Advanced Analytical Technologies, Inc., Ames, Iowa) to assess library size and presence of remaining adapter dimers. This was followed by qPCR assay using KAPA Library Quantification Kit using their SYBR^®^ FAST qPCR Master Mix to estimate the size and quantification. These WGS libraries were sequenced on the Illumina HiSeq-X instrument to generate 150 bp paired-end reads. All flow cell data (BCL files) are converted to barcoded FASTQ files.

### Mapping and variant calling

BWA-MEM version 0.7.12-r1039^31^ was used to align Illumina reads to the owl monkey reference assembly Anan_2.0 (GenBank assembly accession GCA_000952055.2) and to generate BAM files for each of the 30 individuals. Picard MarkDuplicates version 1.105 (http://broadinstitute.github.io/picard/) was used to identify and mark duplicate reads. Single nucleotide variants (SNVs) and small indels (up to 60bp) were called using GATK version 3.3-0 following best practices^32,33^. HaplotypeCaller was used to generate gVCFs for each sample. Joint genotype calling was performed on all samples using GenotypeGVCFs to generate a VCF file. GATK hard filters (SNPs: “QD < 2.0 || FS > 60.0 || MQ < 40.0 || MQRankSum < −12.5 || ReadPosRankSum < −8.0”; Indels: “QD < 2.0 || FS > 200.0 || ReadPosRankSum < −20.0”) (https://software.broadinstitute.org/gatk/documentation/article?id=2806) were applied and calls that failed the filters were removed.

GATK’s PhaseByTransmission was used to identify Mendelian violations that represent possible *de novo* variants. After removing Mendelian violations (MVs) that resulted from missing genotypes or had other anomalies (i.e. 5 MVs with read depth of 0 and 1,984 MVs with allelic depth of 0,0), we obtained 45,432 putative Mendelian violations. We also identified 62 scaffolds as deriving from the X chromosome. These scaffolds had significantly higher homozygosity and lower mean read depth among males (one-tailed t-test, q < 0.05 for both mean homozygosity and read depth). MVs on these scaffolds and scaffolds shorter than 10 kb were removed. This resulted in an initial set of 34,189 putative MVs.

### Filtering of putative mutations

Stringent filters are necessary to avoid potential false positive calls of *de novo* mutations^3,34,35^. To address this issue we applied the following filters to our initial set of MVs:

1. Removed 32,638 MVs with allelic balance less than 0.4 or greater than 0.6 in the child.
2. Removed 112 MVs that are not homozygous reference in both parents.
3. Removed 636 MVs with read depth below 20 or above 60 in any individual in the trio.
4. Removed 520 MVs where the alternate allele is present in an unrelated individual in the sample.

We define allelic balance as the fraction of reads that are a non-reference allele at a given site, meaning that a true heterozygous site should have allelic balance of roughly 0.5. Importantly, we observed that 95% of all initial MVs have allelic balance less than 0.4 (Figure S1a). This indicates that many of these initial calls are false positives. After these four filtering steps we find a total of 283 *de novo* mutations across our 14 trios (Table S2).

### Phasing mutations

Genotypes from three generations allow us to trace the parent of origin for *de novo* mutations transmitted to the third generation. We accomplished this by phasing chromosomal segments with respect to the grandparents (P generation in Fig1a). Phase informative sites were identified in each family and assembled into haplotype blocks. We selected bi-allelic informative sites where: the grandparents had different genotypes, their offspring was heterozygous, and this individual’s partner and offspring were not both heterozygous. The transmission of alleles at these sites can be unambiguously traced to one of the grandparents. We assembled these sites into blocks under the assumption that no more than one recombination occurred per 0.5 Mb interval^20,36^. The phases of haplotype blocks supported by fewer than 100 informative sites were left unassigned, as were the phases of short scaffolds (less than 0.5 Mb). The parent of origin for *de novo* mutations transmitted to the third generation can then be established from the phase of their corresponding haplotype block.

### Estimating mutation rates

To estimate mutation rates per generation per site (*μ_g_*) we must consider rates of error. Our stringent filters ensure that we have few to no false positives; however, we expect that these filters removed a number of true *de novo* variants, leading to a substantial false negative rate (*α*). To estimate *α* resulting from the allelic balance filter, we used the distribution of allelic balance from the total set of 471,532,403 heterozygous autosomal sites in our sample. Unlike the initial set of MVs, the distribution of allelic balance for these sites conforms to the expected distribution for true heterozygous sites, with a single peak at about 0.5 (Figure S1b). We find that the number of heterozygous sites with allelic balance below 0.4 or above 0.6 is 206,358,774 resulting in an estimate of *α* = 0.44. With a less stringent allelic balance filter of 0.3-0.7 the false negative rate falls to 0.29, but changing this filter does not greatly impact the number of mutations called (Figure S2). We correct the observed number of mutations (*m_g_*) in each trio using *α* = 0.44 and an assumed false positive rate of 0. After correction we estimate that there are about 36 *de novo* mutations passed on in a single owl monkey generation.

To calculate the mutation rate per site, we counted the number of callable sites in each trio (*C*). A site was determined to be callable if it passed filters (1) and (4) in the child, filter (2) in the parents, and filter (3) in all individuals in the trio. We find an average number of callable sites of 2,207,614,768 in our 14 trios (range: 2,198,415,883-2,214,425,687). Mutation rates were then calculated by dividing the number of observed mutations (corrected for *α*) in a trio by 2 times the number of callable sites:

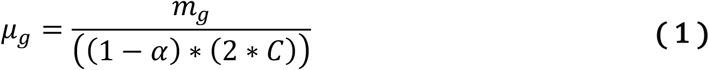

This results in mutation rates ranging from 0.63 × 10^−8^ to 1.5 × 10^−8^ with an average mutation rate of 0.81 × 10^−8^ among the 14 trios (Table S3). Mutation rate was then regressed on father’s age (*A_M_*) (Figure 1, solid line) with the resulting formula for a best fit line:

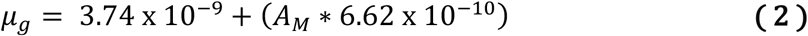

With an average haploid genome size of 2.21 billion base pairs, this means that 19.45 mutations accumulate in males and females before puberty at age 1 and that there are 2.92 additional mutations for every year of the father’s life after puberty in owl monkeys.

### Modeling mutation rates

Large-scale pedigree sequencing projects in humans have shown the importance of different life-stages in the determination of mutation rates^3,11–17,35,37^. Models for predicting mutation rates generally account for the three important life stages in the mammalian germline^19,26^. These life stages are (1) female (*F*), (2) male before puberty (*M*0), (3) and male after puberty (*M*1). The relative contribution of each of these stages must be accounted for when estimating mutation rates per generation^26^ or per year ^19,26,38^. Here, we re-frame this model in terms of reproductive longevity. Reproductive longevity depends on both the age of puberty in males (*P_M_*) and the age of the father at conception of his offspring (*A_M_*) and we find that it is the main determinant of mutation rate variation in primates. We define the value of reproductive longevity (*RL*) as:

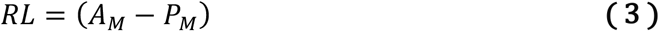

*RL* therefore measures the amount of time mutations have accumulated post-puberty in a male, which only occurs during stage *M*1.

To see how reproductive longevity affects the per-generation mutation rate, *μ_g_*, we must model the combined contribution from all life stages. In any given period of time *t*, the mutation rate due to errors in DNA replication, *μ_t_*, is simply a product of the mutation rate per cell division, *μ_c_*, and the number of cell divisions that occur, *d_t_*:

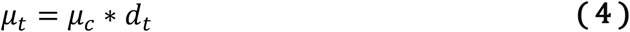

Since females (stage *F*) and pre-puberty males (stage *M*0) have a fixed number of cell divisions, their contribution to the mutation rate per-generation is constant and requires only the substitution of appropriate terms into equation 4:

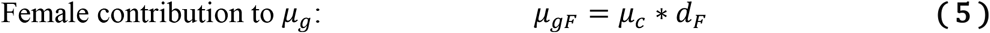

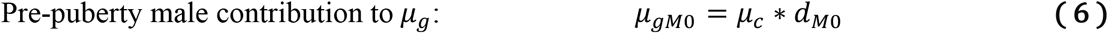

However, in males after puberty (stage *M*1) the number of cell divisions is a linear function of time, and the mutation rate per-generation in this life stage therefore depends on the yearly rate of cell division (*d*_*yM*1_) and reproductive longevity (*RL*):

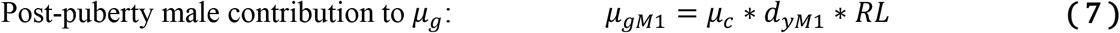

Finally, since an autosome will spend roughly half of its time in females and half in males, the mutation rate per generation (*μ_g_*) for a given species is the average of the male and female contributions:

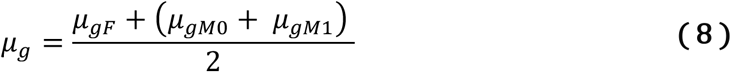

Given estimates of the underlying mutational parameters, this model allows us to predict the mutation rate as a function of reproductive longevity. In order to assess reproductive longevity in species that reach puberty at different times, we used published values for *P_M_*. For owl monkeys, we set *P_M_* at 1 year^21^ (purple line in Figure 3), for humans, we used a value of *P_M_* of 13.4 years^39^ (orange line in Figure 3), for chimpanzees we used 7.5 years^40^ (red line in Figure 3). The ages at conception for all parents in all studies of the mutation rate (points in Fig. 3; Table S4) were taken from the original papers^3,11,12,14–17,20,35,41^.

### Estimating mutational parameters from humans

Empirical observations from developmental studies and large-scale pedigree data from humans inform us about some of the underlying mutational parameters of our model (equations 5, 6, and 7). For example, we use 31 and 34 as estimates for the number of cell divisions in human females (*d_F_*) and males before puberty (*d*_*M*0_)^42^. We use 16 days as the length of a single spermatogenic cycle (*t_sc_*)^43^, which means we expect *d*_*yM*1_ = 23 spermatogenic cycles to occur in a year if all spermatagonial cells are constantly dividing (but see next paragraph).

The remaining parameter of the model, *μ_c_*, can be estimated from human pedigrees. We confirm the estimate of *μ_c_* made by Amster and Sella^38^ by using the *μ_gF_* observed in Kong et al.^3^ of 14.2, the number of female germline divisions, and rearranging equation 5:

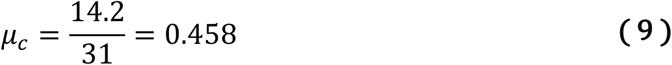

or 1.74 × 10^−10^ given a haploid genome size of 2.63 billion base pairs^3^. We assume this rate is the same between females and males before puberty. However, the observation that 2.01 mutations are passed on per year from the father after puberty^3^ (the mutation rate per year in this lifestage, *μ*_*yM*1_) could imply two things about *μ_c_* in this life-stage: either the mutation rate per cell division has been reduced by an order of magnitude in males after puberty to 0.33 × 10^−11[38]^ or there are fewer than the expected 23 cell divisions per year^44^. There is no evidence to support such a dramatic reduction in the mutation rate per cell division, especially since there does not appear to be a large shift in mutational mechanisms between life stages^17^. The hypothesis that fewer cell divisions have taken place is also more likely based on observations that, of the two types of spermatagonial cells observed in humans, pale and dark, only pale cells actively divide^44,45^. If dividing pale cells transition into non-dividing dark cells and vice versa, then not all spermatagonial cells necessarily undergo 23 spermatogenic cycles in a year and we must re-estimate *d*_*yM*1_. If we assume the mutation rate per cell division in humans is constant before and after puberty, we can estimate the expected number of spermatogenic cycles per year (*d*_*yM*1_):

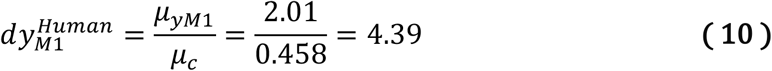

This implies that roughly only 19% of spermatagonial cells are in the pale dividing state at any given time.

Though either a decreased *μ_c_* in males after puberty or a decreased proportion of dividing spermatagonial cells can be fit equally well to the model, we make predictions with the latter sassumption. When predicting a mutation rate function for owl monkeys (Figure 1) we also decrease the length of the spermatogenic cycle to 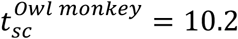 days^46^ and adjust the expected *d*_*yM*1_ assuming 19% of spermatagonial cells are undergoing spermatogenesis at one time:

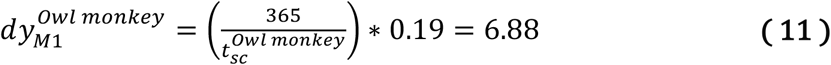

However, when comparing mutation rate functions between species (Figure 3) all underlying mutational parameters are those estimated above from observations in humans, in order to demonstrate that minimal changes to the model can still make accurate predictions of mutation rate functions. Using species-specific parameters of spermatogenesis does not change our results (Fig S3).

## Supplementary figures

**Figure S1:**
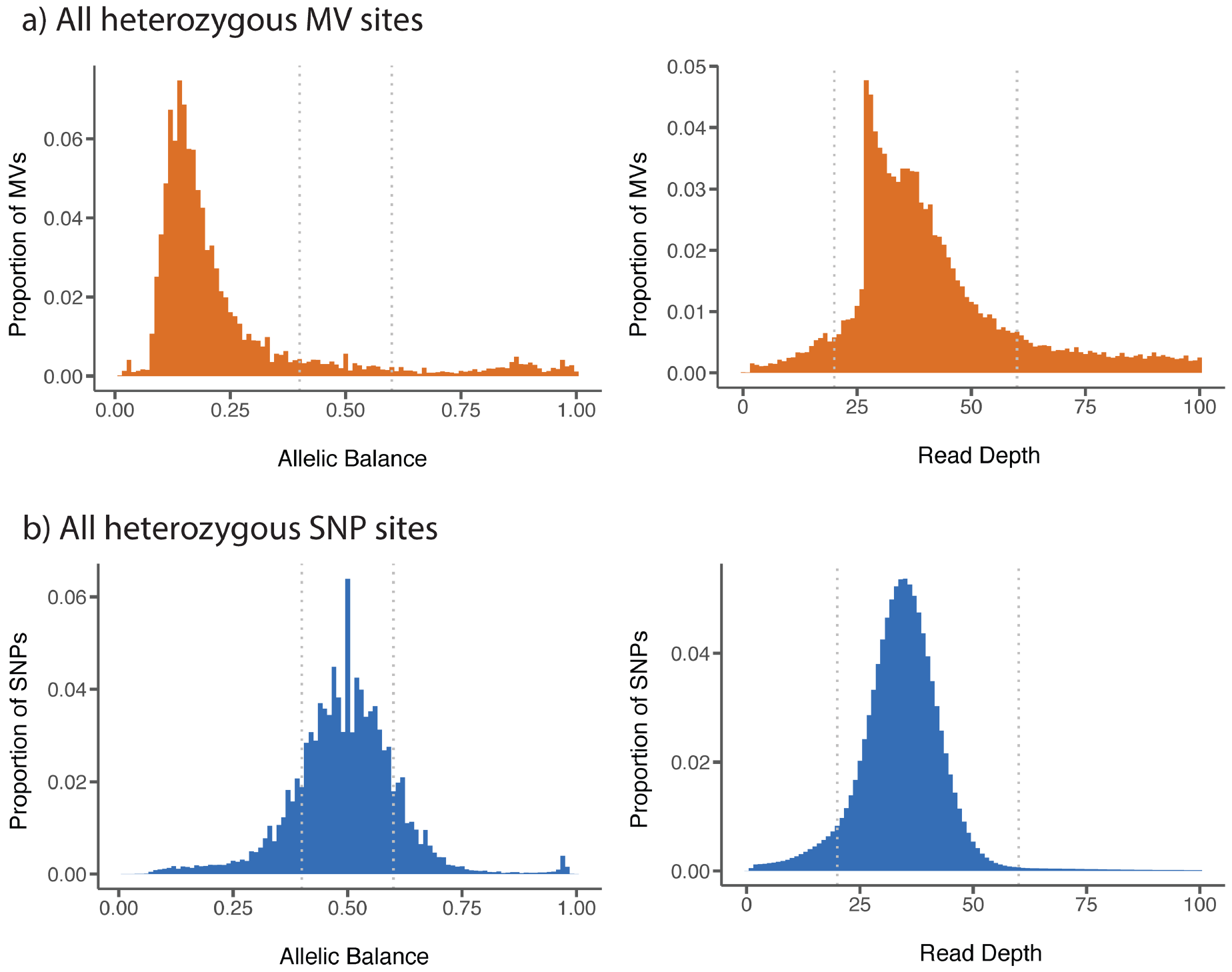
Allelic balance and read depth distributions for **a,** unfiltered Mendelian violations and **b,** all SNP sites in the sample. Grey dotted lines indicate the filtering cutoffs used.

**Figure S2:**
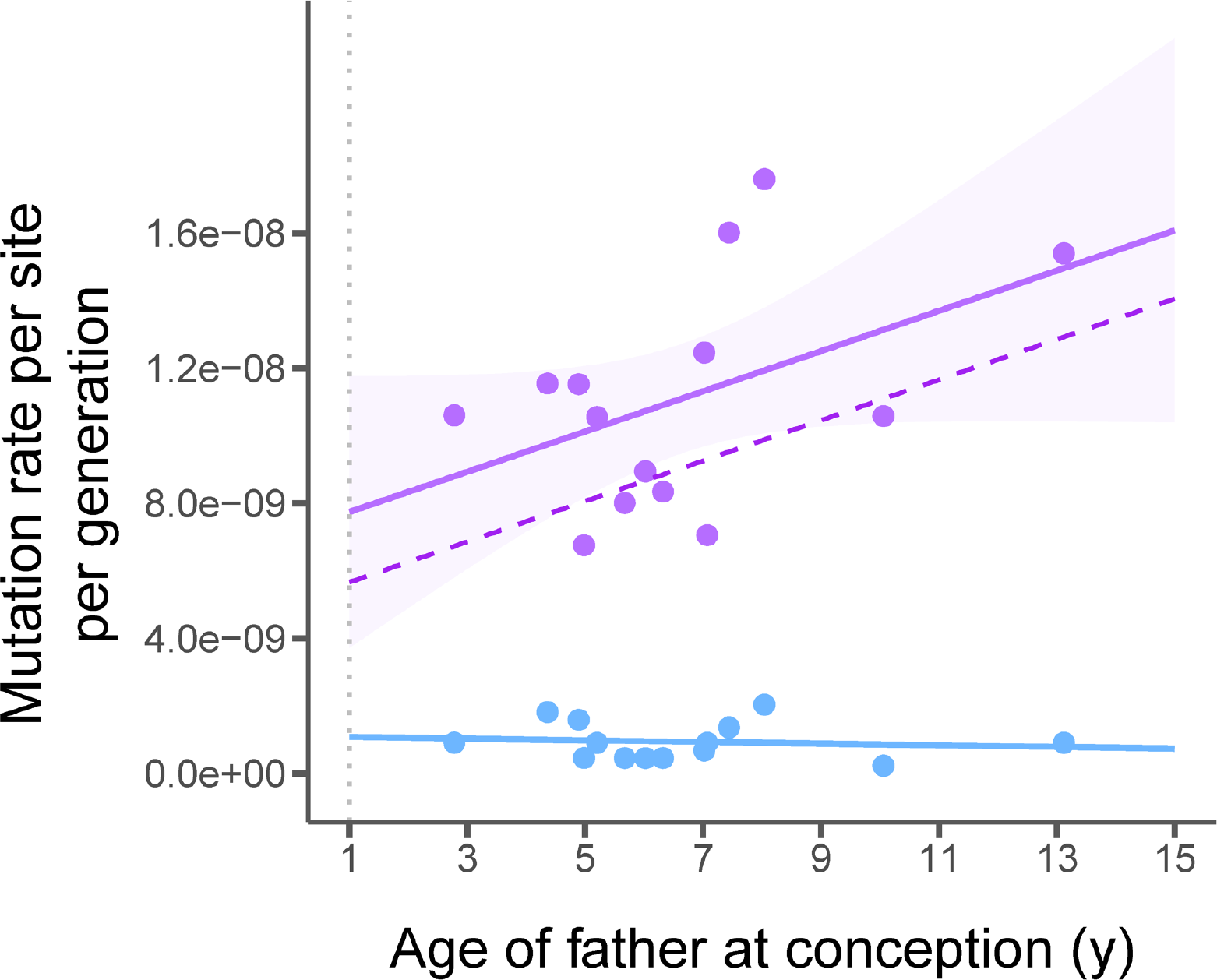
Mutation rates for the 14 owl monkey trios while removing all putative mutations with allelic balance less than 0.3 or greater than 0.7 (purple dots). A linear regression still shows a correlation with father’s age (solid purple line) that is not significantly different from our model’s prediction (dashed purple line; *>F*=1.0, d.f.=13, *P*=1.0). Mutations at CpG sites (blue dots) are not correlated with father’s age (blue line). The grey dotted line indicates age of puberty for owl monkeys.

**Figure S3:**
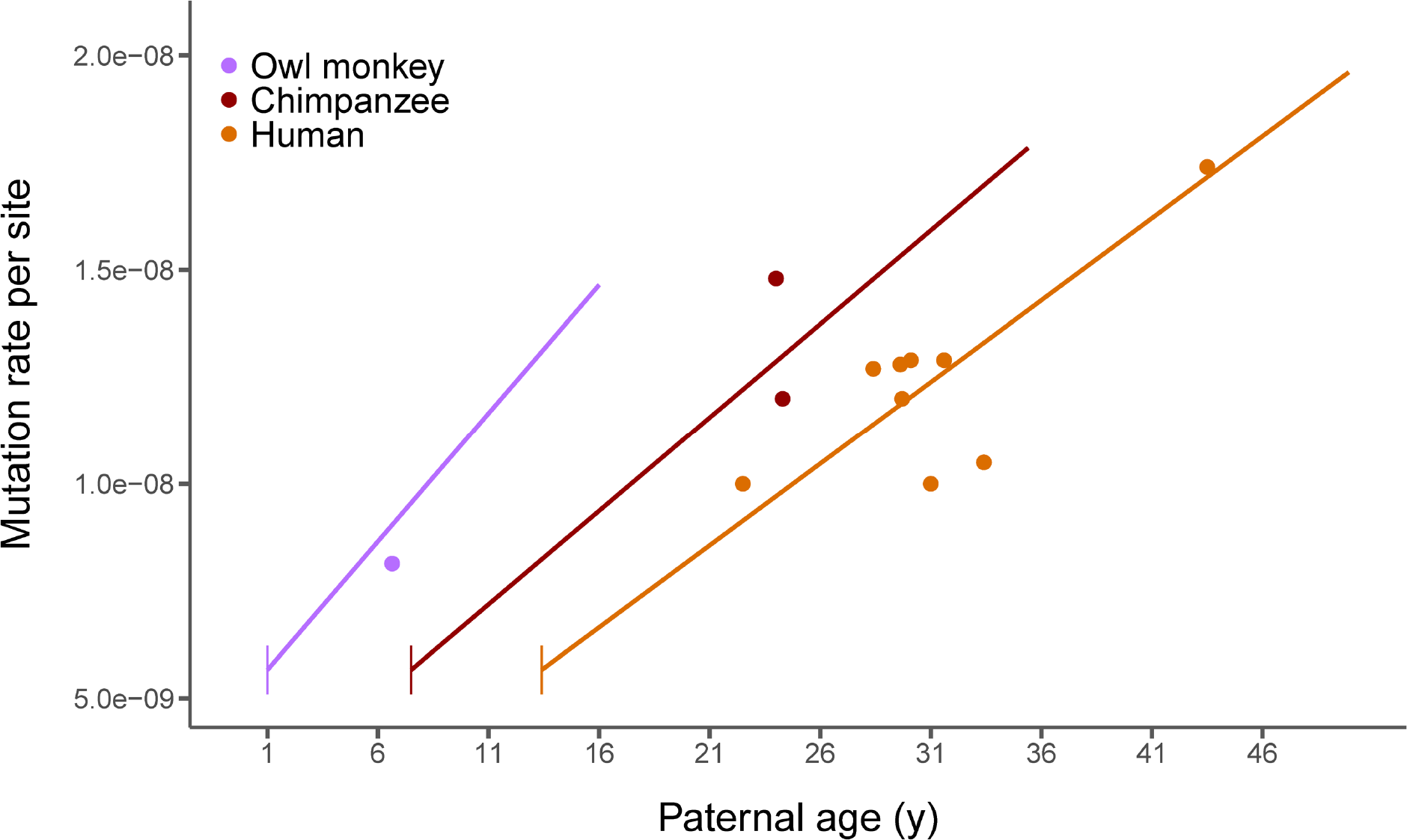
Using species specific rates of spermatogenesis for the three species with high-quality mutation estimates when making predictions from our model of reproductive longevity (lines; equations 3-8 in Methods) does not greatly affect the fit of the model to published estimates of mutation rates (points; see Table S4 for references). Vertical line segments represent the age of puberty for each species. In this figure we used 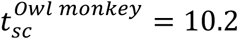(ref. 46), 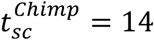 (ref. 47), 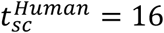 (ref. 43)

